# Synuclein alpha accumulation mediates podocyte injury in Fabry nephropathy

**DOI:** 10.1101/2021.12.17.473164

**Authors:** A Abed, F Braun, D Sellung, M Woidy, O Eikrem, N Wanner, K von Cossel, N Muschol, SW Gersting, AC Muntau, O Kretz, B Najafian, C Tøndel, M Mauer, T Bork, F Grahammer, W Liang, T Eierhoff, W Römer, HP Marti, VG Puelles, C Schell, TB Huber

## Abstract

Current therapies for Fabry disease are based on reversing intra-cellular accumulation of globotriaosylceramide (Gb3) by enzyme replacement therapy (ERT) or chaperone-mediated stabilization of the defective enzyme, thereby alleviating lysosome dysfunction. However, their effect in the reversal of endorgan damage, like kidney injury and chronic kidney disease remains unclear. First, ultrastructural analysis of serial human kidney biopsies showed that longterm use of ERT reduced Gb3 accumulation in podocytes but did not reverse podocyte injury. Then, a CRISPR/CAS9-mediated *α*-Galactosidase knockout podocyte cell line confirmed ERT-mediated reversal of Gb3 accumulation without resolution of lysosomal dysfunction. Transcriptome-based connectivity mapping and SILAC-based quantitative proteomics identified alpha-synuclein (SNCA) accumulation as a key event mediating podocyte injury. Genetic and pharmacological inhibition of SNCA improved lysosomal structure and function in Fabry podocytes, exceeding the benefits of ERT. Together, this work reconceptualizes Fabry-associated cell injury beyond Gb3 accumulation, and introduces SNCA modulation as a potential intervention, especially for patients with Fabry nephropathy.

## Introduction

Anderson-Fabry disease (FD), is an X-linked lysosomal storage disorder caused by mutations in the *GLA* gene (1), which results in an impairment of the hydrolase alpha-Galactosidase A (aGAL). This enzyme deficiency leads to lysosomal dysfunction (2) via progressive accumulation of globotriaosylceramide (Gb3) and other glycosphingolipids (3) in most cells of the body (4). Patients with FD suffer from extensive and progressive end-organ damage, for example cardiomyopathy and nephropathy, both of which key factors for long-term survival (5).

The first therapies became available in 2001 with enzyme replacement therapy (ERT) (6, 7) and were complemented by chaperone therapy in 2016 (8). All have proven efficient in decreasing Gb3 deposits (9–14), but their impact on the reversal of end-organ damage remains unclear.

Glomerular epithelial cells (podocytes) are primary targets in chronic kidney diseases that progress to the requirement of dialysis or transplantation (15–18). Since podocytes exhibit limited regenerative capacities (19, 20), injury and loss are considered critical steps in renal pathophysiology and central therapeutic targets. Previous studies have shown podocytes to accumulate the highest amount of Gb3 in Fabry nephropathy, resulting in early (micro-)albuminuria (21, 22). However, the molecular mechanisms of Fabry podocytopathy remain elusive.

In this study, we report prevailing signs of podocyte damage in human kidney biopsies before and after ERT despite significant Gb3 reduction. We established a CRISPR/Cas9-based GLA knockout in human podocytes, which recapitulates classical cell injury features like lysosomal dysfunction. Quantitative proteomics combined with a network medicine approach identified the accumulation of alpha-Synuclein (SNCA) as a mediator of lysosomal dysfunction resistant to ERT both in vitro and in patient biopsies. This is the first report showing that accumulation of SNCA directly contributes to lysosomal impairment and disease severity in FD in a substrate independent fashion, which suggests that pharmacological targeting of SNCA could serve as an additional therapeutic strategy, especially for patients with Fabry nephropathy.

## Results and discussion

### Podocyte injury persists despite of ERT and significant reductions in Gb3 deposits

Ultrastructural analysis of serial human kidney biopsies (full demographics available in **Table S1**) revealed typical accumulation of Gb3 within podocytes combined with classical signs of podocyte injury, including alterations in foot process morphology (**Fig. 1A**). In agreement with previous reports (13, 14, 23, 24), ERT significantly reduced the volume of podocyte specific Gb3 inclusion bodies in follow-up biopsies. Importantly, increased foot process width remained unaffected (**Fig. 1B**), suggesting that despite significant reductions in Gb3 accumulation and reported correlation of foot process width to Gb3 inclusions (24) remaining Gb3 or independent mechanisms lead to prevailing podocyte injury (25). These observations have direct therapeutic implications, as ERT, if not initiated in the early years of a patient’s live, may not be sufficient to prevent and reverse end-organ damage.

**Figure 1:**
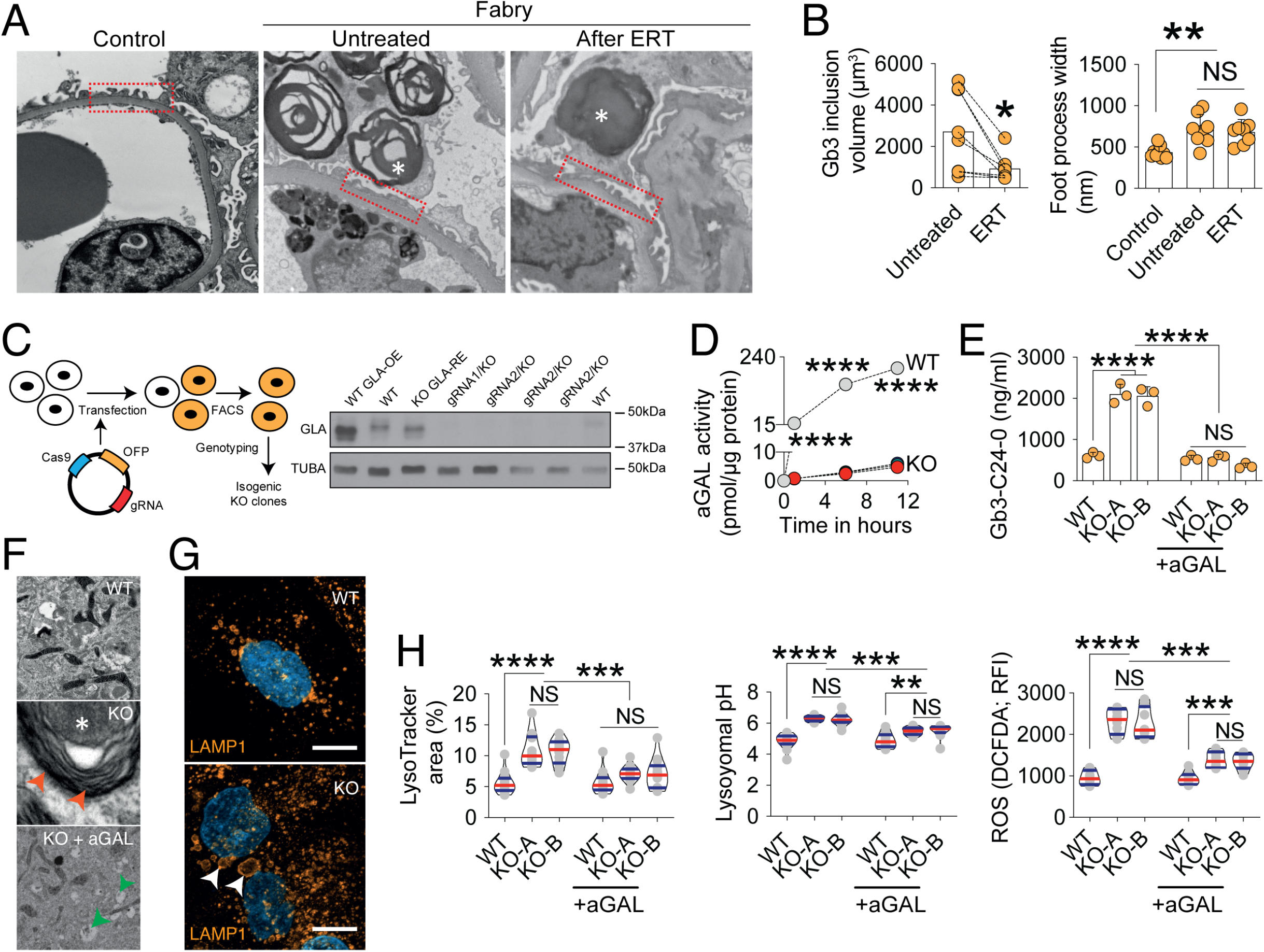
(A) Transmission electron microscopy (TEM) comparison of foot processes between control vs Fabry kidney biopsy before and after ERT with many foot processes widened in both untreated and after ERT Fabry biopsies. Asterisks show Gb3 inclusions in podocytes. (B) Significant decrease of podocyte Gb3 inclusions after ERT but persistence of increased foot process width. (C) Schematic overview of *GLA* knock-out (KO) podocyte generation by CRISPR/Cas9 genome editing. (D) Western blots show a complete absence of GLA expression in several GLA-KO clones. (E) Abolished GLA activity in two KO clones compared to WT cells. (D) Mass spectrometry analysis confirms the accumulation of GB3-C24-0 isoform in KO cells, normalized upon 96 hours of alpha-GAL therapy (n=3). (F) TEM shows zebra bodies exclusively in GLA-KO clones (red arrows). While WT cells depict a normal ultrastructure, aGAL-treated KO cells have remnant vacuoles (green arrows) without zebra bodies. Scalebar represents 1µm. (G) Lysosomal visualization using LAMP1 staining in differentiated WT and KO cells reveals an increased number and size (arrows) of lysosomes in the GLA-KO cells. Scalebars represent 10µm. (H) Quantification of lysosomal area (n=14), pH (n=12) and ROS production (n=8). Violin plots indicating median (red) and upper and lower quartile (blue), **P<0.01, ***P<0.001, ****P<0.0001.

### Generation of a GLA-deficient podocyte line using CRISPR/Cas9 genome editing

The study of end-organ damage in FD has been directly hampered by limited access to human biopsy material. Furthermore, existing animal models do not completely reflect the human phenotype, especially those observed in Fabry nephropathy (26–28).

Thus, we decided to develop a novel in vitro system to model FD-related podocyte damage. Using two independent guide-RNAs (gRNAs), we targeted the first exon within the *GLA* locus to generate *GLA*-deficient immortalized male human podocytes (29) using CRISPR/Cas9 (30, 31) (**Fig. 1C**), eliminating residual intact and functional enzyme (32, 33). Subclones with deletions resulting in premature STOP codons (**Fig. S1A**) and efficient knockout on the protein level (**Fig. 1C**) were selected for further analyses. aGAL activity was almost completely abolished (**Fig. 1D**), resulting in a significant increase of Gb3 in both thin layer chromatography (**Fig. S1B**) and lipid mass spectrometry (**Fig. 1E** and **Fig. S1C-E**). Multilaminar inclusions (zebra bodies) were detected within GLA deficient podocytes with almost complete clearance of these structures after aGAL treatment (**Fig. 1F**).

### In vitro ERT does not fully revert podocyte injury

Both lysosomal number as well as size were dramatically increased in GLA knockout clones and were partially reverted by aGAL treatment (**Fig. 1G** and **Fig. 1H**). This amelioration could not be achieved for lysosomal pH and oxidative stress via reactive oxygen species (ROS) without changes in levels of oxygen consumption rate (OCR) (**Fig. S2A**) or mitochondrial morphology (**Fig. S2B** and **Fig. S2C**). Similar findings have been previously reported in primary fibroblasts derived from Gaucher patients (34) and endothelial cells exposed to Gb3 (35). Furthermore, we detected an increase in autophagy via the decrease of surrogate marker p62, while LC3-II was unchanged both at baseline and in chloroquine challenged KO cells (**Fig. S3**), confirming previous reports (33, 36). Together, our data suggests that podocyte injury extends beyond substrate accumulation, as multiple features of FD are only ameliorated by aGAL replacement.

### SNCA accumulates in GLA-deficient podocytes and is resistant to short-term ERT

We employed SILAC-based quantitative proteomics and transcriptome profiling via RNA sequencing to determine potential alterations in gene expression and protein abundance underlying the observed lysosomal dysfunction (**Fig. S4A**). Filtering for lysosome-associated proteins resulted in the detection of 321 differentially expressed proteins (**Fig. 2A**). Surprisingly, the top-20 regulated lysosomal proteins (except for ANPEP) were not altered at a transcriptomic level (**Fig. 2B**). Interestingly, we observed increased levels of GBA protein, as well as other well described lysosome associated proteins such as LIMP2 (encoded by the gene SCARB2). The latter has been shown to serve as a specific receptor for glucocerebrosidases and to be involved in proper lysosomal biogenesis (37, 38), suggesting that lysosome impairment extends beyond the initial enzyme defect.

**Figure 2:**
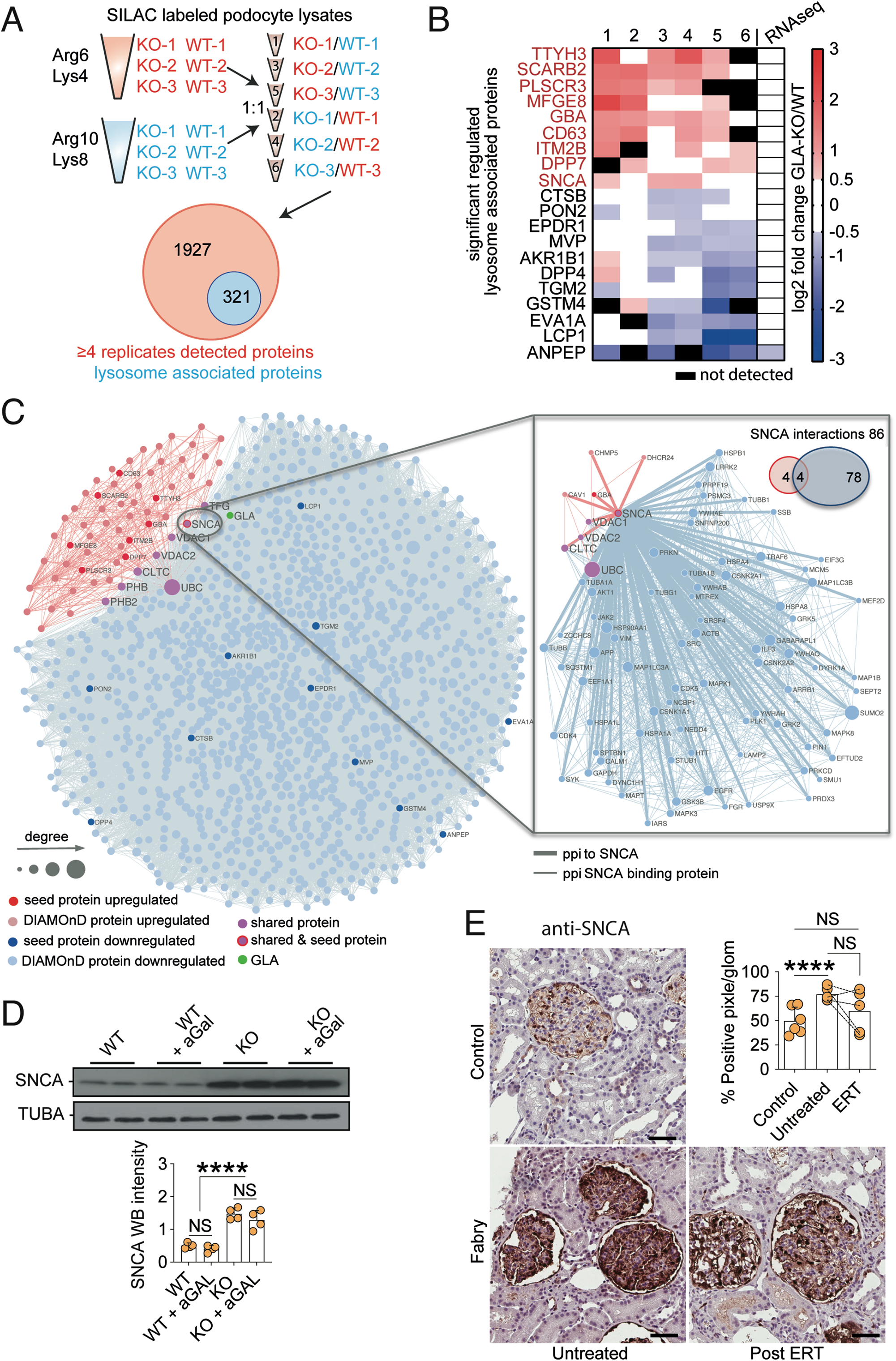
(A) Schematic overview of mass spectrometry analysis using SILAC labelled WT and KO clones. Mass spectrometry yielded 2248 proteins among which 321 proteins are lysosomal enriched. (B) The top ten up and down regulated lysosomal enriched proteins. (C) Network based analysis of up- and downregulated lysosomal proteins associated to GLA knock-out (KO). Nodes represent genes and are connected if there is a known protein interaction between them. The node size is proportional to the number of its connections. Red and blue nodes represent up- and downregulated seed proteins, respectively. Light-red and light-blue nodes represent the respective DIAMOnD proteins. GLA is depicted as green node. Pink nodes indicate shared proteins between the two modules. The separate network and Venn diagram on the right shows the number and interaction partners of SNCA. (D) Western blots of SNCA and TUBA in vehicle and aGAL treated WT and KO cells with quantification confirming the over-expression and resistance of this protein to aGAL treatment (n=4). (E) SNCA staining in representative images and quantification of human renal biopsies showing increase in untreated Fabry samples with resistant accumulation in patients who underwent 5 years of ERT (n=5). Scalebars indicating 50µm. ****P<0,0001

Next, we used a network medicine approach to further evaluate how these proteins are functionally related to each other. We identified the two podocyte-specific GLA KO modules for the respective up- and downregulated lysosomal proteins (**Fig. 2B, S4B-C**). The upregulated hits resulted in a smaller more specific module (64 proteins; z-score: 28). Gene ontology enrichment for cellular compartment and reactome pathways revealed an involvement of processes associated with membrane trafficking, autophagy, mitophagy and the lysosome in both modules (**Fig. S5A-B**). Molecular Function GO terms overrepresented in the disease modules were associated to proteins binding to phosphorylated residues and beta-adrenergic signalling kinases (**Fig. S5C**).

Strikingly, alpha-Synuclein (SNCA) was the only protein found as an upregulated protein by the SILAC-based proteome analysis but also by the network-based approach as a protein residing in the downregulated module, connecting both the upregulated and downregulated disease modules as a seed protein. SNCA is highly produced in many cells and constantly degraded through chaperone-mediated autophagy (39). This protein has been implicated in other lysosomal storage diseases (40) and is well-known in synucleinopathy related neurodegenerative diseases such as Parkinson disease (41, 42). Importantly, the aggregation of pathological SNCA isoforms has been reported as toxic to cells (43–45), and the modulation of SNCA-signalling reverses lysosomal clustering (46), suggesting that SNCA may be an intriguing therapeutic target for FD (47).

SNCA binds to four out of the eight proteins that are connecting both modules with each other and the known interaction partners of SNCA are involved in autophagosome and lysosome function such as GABARAPL1, MAPK1, LAMP2 and SQSTM1 (**Fig. 2C**). Treatment with recombinant aGAL over 96 hours mitigated and normalized the expression levels of all top ten upregulated lysosomal proteins except for SNCA (**Fig. 2D, Fig. S6**). In human renal biopsies, SNCA was detected almost exclusively in the glomerular compartment, showing an increased expression in samples of untreated Fabry patients (**Fig. 2E**). SNCA staining intensity in biopsies of patients who underwent 5 years of ERT was partially reduced (without reaching statistical significance) compared to baseline levels (**Fig. 2E**). Remarkably, we did not observe different SNCA levels in patient derived primary urinary cells and no induction through challenging these cells with globotriaosyl-sphingosine (lyso-Gb3) the main degredation product of Gb3 implicated in the disease’s molecular pathology (**Fig. S7A-B**). Together, these data identify SNCA accumulation as a Gb3-independent event during FD-related podocyte injury. Thus, modulation of SNCA-signalling may provide a novel therapeutic target.

### Modulating SNCA accumulation ameliorates Fabry podocytopathy

To elucidate the effect of alterations in SNCA levels in WT and GLA deficient cells, we performed knockdown and overexpression analyses. We achieved a strong inhibition in both knockout and wild-type cells 48 hours after siRNA transfection (**Fig. 3A**). This inhibition was associated with a significant reduction in lysosomal area, lysosomal pH and ROS accumulation (**Fig. 3B**) without complete reversal to wild-type levels. Next, a reverse overexpression of SNCA (**Fig. 3C**) induced pronounced alterations of lysosomal structure (**Fig. 3D**), and marked increases in lysosomal area, pH and ROS production (**Fig. 3E**), confirming a central role of SNCA-signalling in Gb3-idependent podocyte injury.

**Figure 3:**
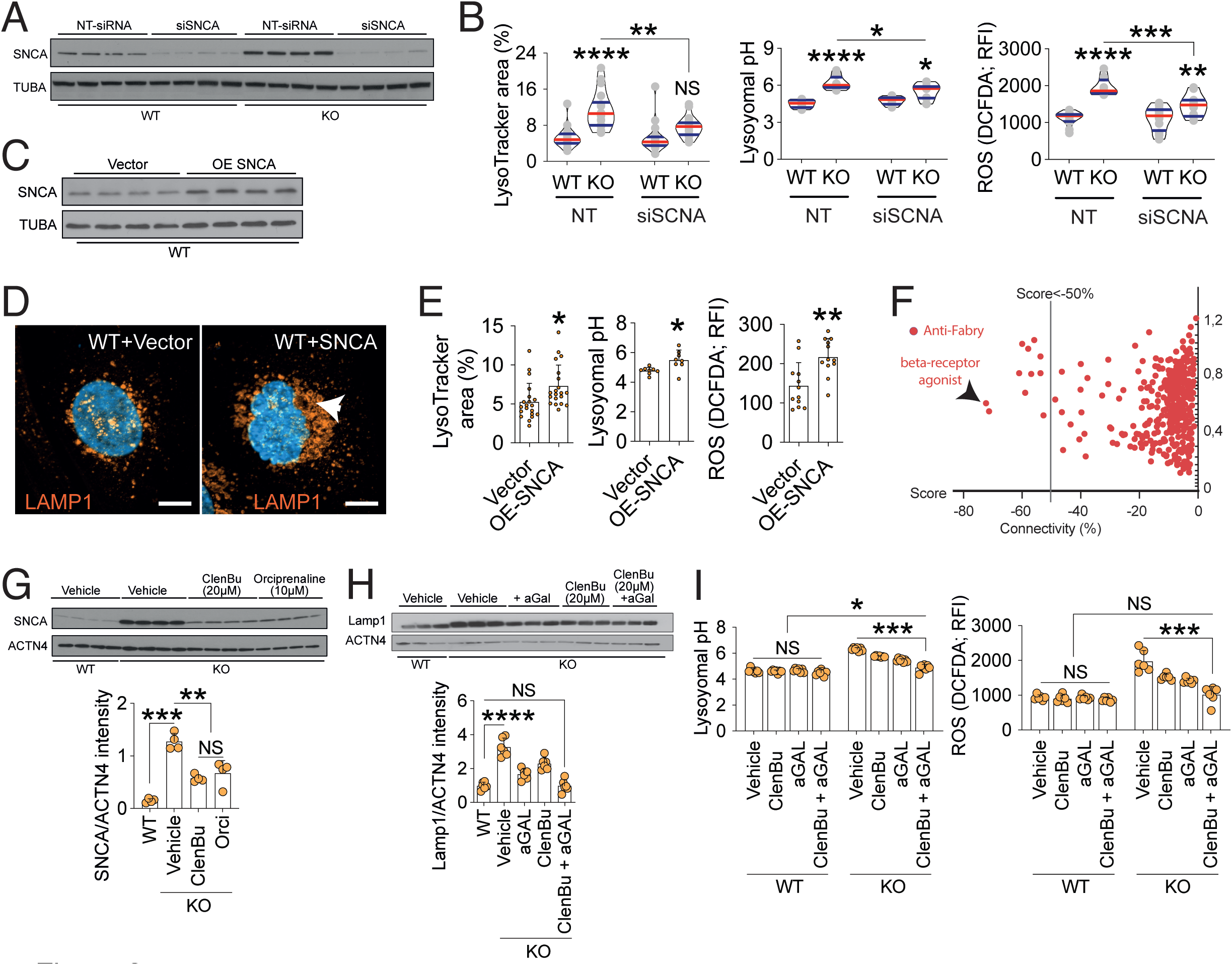
(A) Representative western blot confirming the efficacy of siRNA targeting SNCA in WT and knock-out (KO) clones (n=4). (B) Quantification of lysosomal area, pH and ROS production upon SNCA siRNA treatment (n=18). (C) A representative western blot confirming the overexpression of SNCA in WT cells (n=4). (D) Lamp-1 immunofluorescence staining shows an increase in lysosomal aggregation upon SNCA overexpression (arrow). Scalebars represent 10µm. (E) Quantification of lysosomal area (n=20), pH (n=8) and ROS production (n=12) upon SNCA overexpression. (F) Connectivity mapping showing anti-Fabry compounds with the β2-adrenergic receptor agonist Orciprenaline exhibiting the highest score. (G) Western blots show the expression of SNCA in WT, untreated GLA-KO and KO cells treated with 20µM Clenbuterol and 10µM Orciprenaline (n=6). (H) Western blots depict the expression of Lamp-1 and ACTN4 in WT, untreated GLA-KO and KO cells treated with aGAL, 20µM Clenbuterol, and combined therapy (n=6). (I) Lysosomal pH and ROS analysis in all conditions demonstrate independent and additive effects of β2-adrenergic receptor agonist on lysosomal pH and ROS production in GLA-KO cells (n=6). Violin plots indicate median (red) and upper and lower quartile (blue). Bar graphs depict standard deviation. *P<0.05, **P<0.01, ***P<0.001, ****P <0.0001

### β2 adrenergic receptor agonists as a novel therapy for *Fabry podocytopathy*

Unfortunately, SNCA pharmacological modulators are not currently available. Thus, we performed connectivity mapping analysis of the transcriptomic profile of Fabry podocytes, which identified the β2 adrenergic receptor agonist Orciprenaline to be the top “*anti-Fabry”* compound with a relation score of -0,7 (**Fig. 3F** and **Table S2**). Indeed, Orciprenaline as well as another β2 adrenergic receptor agonist, Clenbuterol, were able to significantly reduce SNCA accumulation in Fabry podocytes (**Fig. 3G**). Furthermore, Clenbuterol depicted a clear dose dependent effect on SNCA (**Fig. S8A-B**). In accordance with the effects of genetic SNCA reduction, β2 adrenergic receptor agonist treatment resulted in decreased lysosomal area (**Fig. S9A**) and increased lysosomal acidification in knockout podocytes (**Fig. S9B**). In line with our findings, it has been shown that beta-2-agonists decrease the risk of Parkinson disease via epigenetic downregulation of SNCA gene transcription and protein reduction (48). Strikingly, the combination of ERT and Clenbuterol showed an additive effect on the restoration of Lamp1 accumulation (**Fig. 3H**) and lysosomal pH and ROS production in Fabry podocytes (**Fig. 3I**), mirroring ultrastructural findings (**Fig. S9C**).

In conclusion, we systematically map the signalling network of Fabry associated kidney disease pathways, identify the role of the accumulation of SNCA contributing to lysosomal impairment and disease severity in Fabry disease and conceptually proof a novel additive pharmacological targeting strategy aiming to halt and reverse Fabry nephropathy.

## Methods

Detailed experimental methods and statistical analyses are included in the Supplemental Methods.

The RNA-Seq and proteome data reported here are deposited in the NCBI’s Gene Expression Omnibus (GEO) database and ProteomeXchange Consortium (GEO GSE179975 & PXD029618).

*Study approval*. Patients in this study provided written informed consent Renal biopsies were performed as part of a clinical trial protocol (ClinicalTrials.gov#: NCT00196716) or standard of care before the initiation of enzyme replacement therapy

## Author contributions

AA, FB & CS designed the research study, conducted experiments, acquired data, analysed data wrote and revised the manuscript. DS, OE, KC, MW, SWG, OK, BN, CT, MM, TB, FG, WL, TE, WR conducted experiments, acquired data and analysed data. OE conducted experiments, acquired data and analysed data. NM provided resources and acquired patient samples. ACM provided resources and revised the manuscript. HPM designed the research study and provided resources. VGP designed the research study, provided resources, wrote and revised the manuscript. TBH designed the research study, provided resources, wrote and revised the manuscript.

## Acknowledgements

We thank Charlotte Meyer, Betina Kiefer and Valerie Oberüber for expert technical assistance. We thank Prof. Em. Einar Svarstad for taking part of the Fabry patient. We would also like to express our gratitude to all members of our laboratories and to the Life Imaging Center (LIC) of the University of Freiburg for helpful discussions and support.

## References

1. Anderson W. A CASE OF “ANGEIO-KERATOMA.”*. British Journal of Dermatology 1898;10(4):113–117.

2. Kuech E-M, Brogden G, Naim HY. Alterations in membrane trafficking and pathophysiological implications in lysosomal storage disorders. Biochimie 2016;130:152–162.

3. Guce AI et al. Catalytic mechanism of human alpha-galactosidase.. J Biological Chem 2009;285(6):3625–32.

4. Schiffmann R. Fabry disease.. Handbook of clinical neurology 2015;132:231–248.

5. Mehta A et al. Natural course of Fabry disease: changing pattern of causes of death in FOS - Fabry Outcome Survey.. J Med Genet 2009;46(8):548–52.

6. Schiffmann R et al. Enzyme Replacement Therapy in Fabry Disease. Jama 2001;285(21):2743.

7. Eng CM et al. Safety and efficacy of recombinant human alpha-galactosidase A--replacement therapy in Fabry’s disease.. The New England journal of medicine 2001;345(1):9–16.

8. Germain DP et al. Treatment of Fabry’s Disease with the Pharmacologic Chaperone Migalastat.. The New England journal of medicine 2016;375(6):545–555.

9. Germain DP et al. Ten-year outcome of enzyme replacement therapy with agalsidase beta in patients with Fabry disease.. Journal of medical genetics [published online ahead of print: March 20, 2015]; doi:10.1136/jmedgenet-2014-102797

10. Mehta A et al. Enzyme replacement therapy with agalsidase alfa in patients with Fabry’s disease: an analysis of registry data.. Lancet 2009;374(9706):1986–1996.

11. Hughes DA et al. Oral pharmacological chaperone migalastat compared with enzyme replacement therapy in Fabry disease: 18-month results from the randomised phase III ATTRACT study.. Journal of medical genetics 2016;jmedgenet-2016-104178.

12. Mauer M et al. Reduction of podocyte globotriaosylceramide content in adult male patients with Fabry disease with amenable GLA mutations following 6 months of migalastat treatment.. Journal of medical genetics 2017;54(11):781–786.

13. Thurberg BL et al. Globotriaosylceramide accumulation in the Fabry kidney is cleared from multiple cell types after enzyme replacement therapy.. Kidney international 2002;62(6):1933–1946.

14. Tøndel C et al. Agalsidase benefits renal histology in young patients with Fabry disease.. Journal of the American Society of Nephrology : JASN 2013;24(1):137–148.

15. Gödel M et al. Role of mTOR in podocyte function and diabetic nephropathy in humans and mice.. The Journal of clinical investigation 2011;121(6):2197–2209.

16. Rinschen MM et al. A Multi-layered Quantitative In Vivo Expression Atlas of the Podocyte Unravels Kidney Disease Candidate Genes.. Cell reports 2018;23(8):2495–2508.

17. Puelles VG et al. mTOR-mediated podocyte hypertrophy regulates glomerular integrity in mice and humans. Jci Insight 2019;4(18):e99271.

18. Kopp JB et al. Podocytopathies. Nat Rev Dis Primers 2020;6(1):68.

19. Wanner N et al. Unraveling the role of podocyte turnover in glomerular aging and injury.. Journal of the American Society of Nephrology : JASN 2014;25(4):707–716.

20. Berger K et al. The Regenerative Potential of Parietal Epithelial Cells in Adult Mice.. Journal of the American Society of Nephrology : JASN [published online ahead of print: January 9, 2014]; doi:10.1681/asn.2013050481

21. Najafian B et al. Progressive podocyte injury and globotriaosylceramide (GL-3) accumulation in young patients with Fabry disease. Kidney international 2010;79(6):663–670.

22. Tøndel C, Bostad L, Hirth A, Svarstad E. Renal biopsy findings in children and adolescents with Fabry disease and minimal albuminuria.. American journal of kidney diseases : the official journal of the National Kidney Foundation 2008;51(5):767–776.

23. Najafian B et al. Accumulation of Globotriaosylceramide in Podocytes in Fabry Nephropathy Is Associated with Progressive Podocyte Loss.. J Am Soc Nephrol Jasn 2020;ASN.2019050497.

24. Najafian B et al. One Year of Enzyme Replacement Therapy Reduces Globotriaosylceramide Inclusions in Podocytes in Male Adult Patients with Fabry Disease.. PloS one 2016;11(4):e0152812.

25. Braun F et al. Enzyme Replacement Therapy Clears Gb3 Deposits from a Podocyte Cell Culture Model of Fabry Disease but Fails to Restore Altered Cellular Signaling. Cell Physiol Biochem 2019;52(5):1139–1150.

26. Miller JJ et al. α-Galactosidase A-deficient rats accumulate glycosphingolipids and develop cardiorenal phenotypes of Fabry disease.. FASEB journal : official publication of the Federation of American Societies for Experimental Biology 2018;fj201800771R.

27. Ohshima T et al. alpha-Galactosidase A deficient mice: a model of Fabry disease.. Proc National Acad Sci 1997;94(6):2540–2544.

28. Taguchi A et al. A symptomatic Fabry disease mouse model generated by inducing globotriaosylceramide synthesis.. The Biochemical journal 2013;456(3):373–383.

29. Saleem MA et al. A conditionally immortalized human podocyte cell line demonstrating nephrin and podocin expression. [Internet]. Journal of the American Society of Nephrology : JASN 2002;13(3):630–638.

30. Jinek M et al. A Programmable Dual-RNA-Guided DNA Endonuclease in Adaptive Bacterial Immunity. Science 2012;337(6096):816–821.

31. Ran FA et al. Genome engineering using the CRISPR-Cas9 system. Nature protocols 2013;8(11):2281–2308.

32. Labilloy A et al. Altered dynamics of a lipid raft associated protein in a kidney model of Fabry disease.. Molecular Genetics and Metabolism [published online ahead of print: October 19, 2013]; doi:10.1016/j.ymgme.2013.10.010

33. Liebau MC et al. Dysregulated Autophagy Contributes to Podocyte Damage in Fabry’s Disease. Plos One 2013;8(5):e63506.

34. Deganuto M et al. Altered intracellular redox status in Gaucher disease fibroblasts and impairment of adaptive response against oxidative stress. J Cell Physiol 2007;212(1):223–235.

35. Shen J-S et al. Globotriaosylceramide induces oxidative stress and up-regulates cell adhesion molecule expression in Fabry disease endothelial cells.. Mol Genet Metab 2008;95(3):163–8.

36. Chévrier M et al. Autophagosome maturation is impaired in Fabry disease. 2010;6(5):589–599.

37. Gonzalez A, Valeiras M, Sidransky E, Tayebi N. Lysosomal integral membrane protein-2: a new player in lysosome-related pathology.. Mol Genet Metab 2013;111(2):84–91.

38. Slaats GG et al. Urine-derived cells: a promising diagnostic tool in Fabry disease patients. Sci Rep-uk 2018;8(1):11042.

39. Vogiatzi T, Xilouri M, Vekrellis K, Stefanis L. Wild type alpha-synuclein is degraded by chaperone-mediated autophagy and macroautophagy in neuronal cells.. J Biological Chem 2008;283(35):23542–56.

40. Magalhaes J et al. Autophagic lysosome reformation dysfunction in glucocerebrosidase deficient cells: relevance to Parkinson disease. Human molecular genetics 2016;25(16):3432–3445.

41. Picillo M et al. Parkinsonism due to A53E α-synuclein gene mutation: Clinical, genetic, epigenetic, and biochemical features. Movement Disord 2018;33(12):1950–1955.

42. Rosborough K, Patel N, Kalia LV. α-Synuclein and Parkinsonism: Updates and Future Perspectives. Curr Neurol Neurosci 2017;17(4):31.

43. Dehay B et al. Targeting α-synuclein for treatment of Parkinson’s disease: mechanistic and therapeutic considerations. Lancet Neurology 2015;14(8):855–866.

44. Gitler AD et al. The Parkinson’s disease protein -synuclein disrupts cellular Rab homeostasis. Proc National Acad Sci 2007;105(1):145–150.

45. Martin LJ et al. Parkinson’s Disease -Synuclein Transgenic Mice Develop Neuronal Mitochondrial Degeneration and Cell Death. J Neurosci 2006;26(1):41–50.

46. Adams J et al. Autophagy-lysosome pathway alterations and alpha-synuclein up-regulation in the subtype of neuronal ceroid lipofuscinosis, CLN5 disease.. Sci Rep-uk 2019;9(1):151.

47. Wise AH et al. Parkinson’s disease prevalence in Fabry disease: A survey study.. Molecular genetics and metabolism reports 2018;14:27–30.

48. Mittal S et al. β2-Adrenoreceptor is a regulator of the α-synuclein gene driving risk of Parkinson’s disease. Science 2017;357(6354):891–898.

