## Supplementary Information for "Synuclein alpha accumulation mediates podocyte injury in Fabry nephropathy"

### Detailed Materials and Methods

*Quantification of Gb3 deposition and foot process width in renal biopsies*

Subjects consisted of eight (M/F=7/1) patients with classic Fabry disease. Fabry disease was confirmed by measurement of leukocyte alpha-galactosidase A activity and/or GLA sequencing. Studies were performed in accordance with principles of the Declaration of Helsinki and were approved by the Institutional Review Board of the University of Washington, University of Minnesota, and the Regional Ethics Committee of Western Norway. Written informed consent had been obtained from each subject.

Renal biopsies were performed as part of a clinical trial protocol (ClinicalTrials.gov#: NCT00196716) or standard of care before the initiation of enzyme replacement therapy and 11 (n=6) or 12 (n=2) months following treatment with agalsidase-beta (1 mg/kg/every other week). Kidney biopsies from none living transplant donors obtained before organ removal were studied as controls. Semi-thin sections of 2.5% glutaraldehyde-fixed, plastic embedded tissues were stained with toluidine blue for identification of glomeruli. Random glomerular sections were prepared for stereological studies as described elsewhere (1). Overlapping digital low-magnification (~8000X) images of entire glomerular profiles and high-magnification (~30,000X) images of glomeruli according to a systematic uniform random sampling protocol were obtained using a JEOL 1010 electron microscope (2). Volume of GL-3 inclusions per podocyte [V(Inc/PC)] was estimated using a combination of point counting and point-sampled intercept method (1, 3). Podocyte average foot process width (FPW) was estimated as the reciprocal of slit-length density as previously described (4).

*Cell culture*

Conditionally immortalized human podocytes were kindly provided by M. Saleem (University of Bristol, UK). Cells were let to proliferate at 33°C in RPMI-1640 medium supplemented with 10% fetal calf serum (FCS), Penicillin/Streptomycin, ITS and non-essential amino acids. To induce differentiation, 70% confluent podocytes were switched to 37°C for 10 to 14 days (unless indicated otherwise). Cells were let to differentiate over collagen IV coated plates (50 ng/µl, Sigma, Germany) in all experiments. Rescue experiments were performed by adding 20µg/ml alpha-galactosidase-A (Genzyme-Sanofi) to the complete medium. aGAL treatment was applied on 10 days differentiated cells over 96 hours with 24 hours' intervals of enzyme renewal. The $\beta_2$ adrenergic receptor agonist Clenbuterol hydrochloride (European Pharmacopoeia) and Orciprenaline (Sigma Aldrich) were used in our experiments. Different concentrations were applied over 96 hours (as indicated in the figure legends). Likewise, alpha-GAL, $\beta_2$ adrenergic receptor agonist treatment was applied on 10 days differentiated cells and renewed every 24 hours over 96 hours. Primary urinary cells were collected from Fabry patients in the International Center for Lysosomal Storage Diseases (ICLD) Hamburg according to Ethics Statement PV3501 approved by the Ethics Board of the Board of Physicians Hamburg. Cells were taken into culture and expanded as described previously (5, 6). After reaching a sufficient confluency, cells were treated with normal proliferation medium (5) supplemented with DMSO (Vehicle) or 100 nM lyso-Gb3 (Sigma Aldrich) in DMSO for 48h.

*Generation of isogenetic GLA knockout human podocytes*

Crispr/CAS9 genome editing was applied to generate GLA knockout podocytes *in vitro* as previously described (7) . Briefly, Crispr/CAS9 genome editing with two different gRNAs (guideRNA1: 5′-TTGTCCAGTGCTCTAGCCCC (AGG)-3′, guideRNA2: 5′-CAGTGCAGCCAGCCCATGGT (AGG)-3′ targeting the first exon of the human GLA gene was used. A web-based platform was operated in order to design these gRNAs (e-crisp.org - http://www.e-crisp.org/E-CRISP/). The gRNAs being subcloned in targeting CRISPR nuclease vectors with OFP (GeneArt life technologies, Carlsbad, CA, USA) according to the manufacturer's protocol were inserted in the immortalized human podocytes via electroporation. Mixed cell populations were validated with restriction enzyme digestion with NcoI (5'-C||CATGG-3', New England BioLabs Inc., Ipswich, MA, USA) for gRNA2 or with an indel mutations detecting and cleaving the enzyme (Genomic Cleavage Detection Kit, GeneArt life technologies, Carlsbad, CA, USA) for gRNA1. After an incubation period of 48 hours, single cells were selected via FACS, sorted into 96-well plates, and further expanded into isogenetic clone colonies. The DNA sequences were analysed using the Sanger technique after initial DNA isolation and amplification (forward primer: 5′-TGGAAATAGGGCGGGTCAAT-3′, reverse primer: 5′-TTCCCCAAACACACCCAAAC-3′). The sequencing results proved the sex of the clone cell line used to be male. Hemizygous GLA- clones translated in silico were reviewed for frameshift mutations and premature stop codons.

*Electron microscopy of cell culture podocytes*

Cell samples where fixed in 4% paraformaldehyde plus 1% glutaraldehyde in 0.1M phosphate buffer over night After contrastation using 1% osmium tetroxid in 0.1 M phosphate buffer (45 min at RT) and 1% uranyl acetate (in 70% ethanol, RT) samples where dehydrated in an ascending ethanol series and embedded in epoxy resin (Durcupan, Sigma Aldrich). Ultrathin sections of approx. 70nm thickness where prepared using a Leica Ultracut UC6. For imaging Philipps CM100 transmission electron microscope was used.

*RNA sequencing*

Conditionally immortalized human podocyte WT and CRISPR/CAS9 GLA KO cell lines were differentiated at 37°C at 70% confluency for 10 to 14 days. Total RNA of cells was isolated using the Phenol/Chloroform method as previously described (8).

Library preparation and sequencing was performed by GATC, Germany. All raw data was deposited to Gene Expression Omnibus (GEO), accession number GSE186258.

*Connectivity mapping*

The top 100 up and downregulated genes from RNA Sequencing data were uploaded to the next generation connectivity mapping tool (CLUE website) (9). Output analysis and ranking were automatically performed by the website.

*Proteomics analysis*

In order to identify new regulators that may play a role in the lysosomal dysfunction we studied the whole proteome in GLA-KO cells by employing quantitative, SILAC-based proteomics and our novel podocyte Fabry model and by analysing 3 Fabry clones versus 3 WT clones. For MS analysis, SILAC labeling of human immortalized podocytes was performed for 14 days as previously described (Schell et al., 2017). Based on SILAC labeling and protein concentration WT and KO samples were mixed 1:1 for MS-analysis. LC-MS/MS data analysis was performed as reported before (Weise et al., Mol Neurobiol, 2019). The MS raw data files were uploaded into the MaxQuant software version 1.4.1. which performs peak and SILAC-pair detection, generates peak lists of mass error corrected peptides and data base searches. We identified nearly 2,300 proteins among which 321 are lysosomal enriched proteins. The top 10 up and downregulated were considered as targets of high interest for further functional analysis. All raw data and original result files were deposited to the ProteomeXchange Consortium (http://proteomecentral.proteomexchange.org) via the PRIDE partner repository with the dataset identifier PXD029618.

*Building GLA specific network modules*

Network-based approaches are based on the assumption that proteins that participate in the same biological processes or share molecular functions are not scattered randomly but tend to cluster and build functionally relevant modules within the human interactome. In order to identify the *disease module* that connects the detected up- and downregulated proteins associated with GLA knockout, we used a large dataset of known protein interactions recently compiled by Cheng and colleagues (10). It consists of 16,677 proteins (nodes) connected by 243,603 protein interactions (edges). One way to build the module would be to search for known direct interaction partners of the up- and downregulated proteins (seed proteins), calculate the size of the resulting largest connected module between them and evaluate whether this size significantly differs from random expectation. However, this method would favour highly connected proteins in the human interactome. In order to detect connections between the seed proteins in an unbiased way, we used a recently published disease module detection algorithm (DIAMOnD). DIAMOnD takes into account the *connectivity significance* of proteins to a set of seed proteins and helps to build a module around them (11) The algorithm calculates the probability that a protein with k links has exactly $k_s$ links to the seed proteins using the hypergeometric distribution and determines the p-value that it has more connections to the seed proteins than expected. The protein with the lowest p-value is then added to the module and a new iteration starts. Since the algorithm could iterate over all 16,677 proteins of the interactome a break-off criterion to determine the final optimal module size needs to be defined.

*Determining the final module size*

In order to determine the final module size we used the method that was recently described by Halu and colleagues (12). DIAMOnD ranks the proteins that are not seed proteins according to their p-value (see above) and incorporates the protein with the lowest p-value into the module. After each iteration the resulting module size, e.g. the number of nodes that are directly connected to each other is determined, and compared against random expectation (1000 random network samples), resulting in a respective z-score:

$$z - score = \frac{module - randommodule}{\sigma random},$$

where $module$ and $randommodule$ are the sizes of the resulting largest connected module and the random expectation. The standard deviation of the calculated 1000 random module sizes is depicted by $\sigma random$. After each iteration, we determined how many seed proteins are integrated into the module. As proposed by Halu et al. the module size where all seed proteins are integrated into the module and where the corresponding z-score is above 1.96 (significant z-score) is taken as the final module size.

*aGAL enzyme activity measurement*

The fluorometric measuring method of the enzyme activity of alpha-galactosidase-A with 4-Methylumbelliferyl-α-D-galactopyranoside has been previously fully described (13–15). In addition, N-acetylgalactosamine has been shown to significantly inhibit alpha-galactosidase B activity (15). Briefly, cells from several aGLA- KO isogenetic clones and wildtype cells were counted and pelleted. The pellets were lysed in 500µl lysis buffer (27 mmol/l sodium citrate, 46mmol/l sodium phosphate dibasic, 0,1% Triton X-100, 1 M HCL, in ddH2O, pH 4,6) by pipetting on ice. Proteins were then separated by centrifuging at 13,2 g and the protein concentration was determined via the Pierce BCA protein assay. 10µl of each sample with three technical replicates were incubated with 25µl test buffer (27 mmol/l sodium citrate, 46 mmol/l sodium phosphate dibasic, 6 mmol/l 4-Methylumbelliferyl α-D-galactopyranoside, 90 mmol/l N-Acetyl-D-galactosamine, 1 M HCL, in ddH2O, pH 4,6) at 37°C for one, six or eleven hours. Subsequently, 35 µl stop buffer (0,4 mol/l Glycine, 5 N NaOH, in ddH2O, pH 10,8) was added and the fluorescence measured. After initial shaking with amplitude of 1 mm for 10 seconds and a 25-time excitation with 355nm the emission of 455nm were measured over an integration period of 20 µs. A standard activity curve was established using aGAL from green coffee beans (Sigma-Aldrich, Saint Louis, MO, USA) with known aGAL activity in serial dilution.

*Mass-spectrometric measurement of Gb3*

Differentiate WT and KO podocytes were treated 48 hours with alpha-GAL or PBS for Mass-spectrometric Gb3 quantification as described previously (16). By the end of incubation, cells were washed twice with PBS, harvested by trypsin and washed again with ice-cold PBS and directly lysed in 0.2% Triton-PBS. Equal volume was taken from each sample for total protein measurement in order to normalize the final readings (Pierce BCA protein assay). Lysate was then stored at -20 and GB3 quantification were performed by pharm-analyt Labor GmbH, Austria.

*Dual-emission ratiometric measurement of lysosomal pH and Lysotracker*

LysoSensor™ Yellow/Blue DND-160 (Life Technologies) was used according to the manufacturer description in order to determine the lysosomal acidity in cultured cells. Briefly, cells were suspended and labelled with 10 µM LysoSensor DND-160 for 2 hours at 37°C in 10% medium, cells were then pelleted and washed in PBS twice. The labelled cells were treated for 10 min with 10 mM monensin (Sigma) and 10 M nigericin (Sigma) in 25 mM 2- (N-morpholino) ethanesulfonic acid (MES) calibration buffer, pH 3.5–8.0, containing 5 mM NaCl, 115 mM KCl and 1.2 mM MgSO4. Cells were then distributed in black 96-well plate (2500cells/well) and the fluorescence was measured with a TECAN plate reader. Light emitted at 440 and 535 nm in response to excitation at 340 and 380 nm were measured, respectively. The ratio of light emitted with 340 and 380 nm excitation was plotted against the pH values in MES buffer, and the pH calibration curve for the fluorescence probe was generated (17). In order to study the lysosomal structure and to score the cellular lysosomal mass, Lysotracker™ Red DND-99 (Life Technologies) was used according to the manufacturer's recommendation. Lysotracker was applied to differentiated podocytes for 30min at 37°C with a final working concentration of 50 nM. Cells were washed and images were taken using a Zeiss Axio Observer microscope, equipped with a 63x objective and Apotome function and analysed using cell profiler. The Lysosomal fraction was blotted as a percentage of lysosomal to cellular area.

*Oxidative Stress Detection*

DCFDA-Assay (ThermoFisher Scientific) was employed to measure ROS level in adherent cells (18). Differentiated cells were harvested and seeded back in a dark, clear bottom 96-well microplate with a concentration of 2500 cells per well and allowed to adhere overnight. Cells were then washed with PBS and incubated with DCFDA solution (20 µMol in PBS) by adding 100 µl/well for 45 minutes at 37°C in the dark. DCFDA Solution was removed, and wells were washed twice with PBS and the florescence was immediately read by TECAN reader (Ex/Em= 485/535 nm).

*Thin layer chromatography*

Lipid extraction was carried out in principle according to the method of Bligh and Dyer (19). Pelleted cells were lysed by osmotic shock using ddH2O, afterwards transferred into a glass tube, mixed and vortexed with chloroform:methanol (1:2) for 1 min. Then, 1.25 mL of chloroform and 1.25 mL of ddH2O was added and mixed for 15 sec. For phase separation, the solution was centrifuged at 3000 rpm for 10 min at 15°C. The organic phase was transferred to a new glass tube. The hydroalcoholic phase was washed once with 1.5 mL of chloroform, mixed for 15 sec and then centrifuged at 4000 rpm for 10 min at 15°C. The organic phases were combined and the solvent was evaporated under a stream of nitrogen. Dried lipids were either stored under an argon atmosphere or resolved in chloroform:methanol (2:1) for thin-layer chromatography (TLC). Lipid extracts were loaded on two silica plates and separated by TLC using methanol:chloroform:ddH2O (60:35:8) as a mobile phase. Afterwards one silica plate was incubated with p-anisaldehyde/acidic alcohol solution and placed in an oven for 15 - 30 min at 120°C to stain overall lipids. For specific detection of Gb3, the other plate was fixed by 3.75 % (w/v) polyisobutylmethacrylate/n-Hexane, blocked with 1% (w/v) BSA/PBS (containing calcium and magnesium), incubated afterwards with biotinylated Shiga toxin B-subunit (1.8 µg/mL) and subsequently with alkaline phosphatase-conjugated streptavidin (2 µg/ml). Gb3 bands were visualized by a colorimetric reaction with NBT/BCIP Substrate Solution (Thermo Scientific). The chromatogram was densitometrical analysed and documented using Fusion FX (Vilber Lourmat) and ImageJ software.

*Seahorse XFp mitochondrial analysis and ATP measurement*

Optimization of cell density for human podocytes of the respective genotype as well as optimization of the working concentration titers for each individual inhibitor was conducted prior to the Seahorse XFp experiments according to the manufacturer`s instructions (Agilent Technologies). Human podocytes were seeded at a density of 15 000 cells/well. The Seahorse XFp Mito Stress Test was performed following the manufacturer`s instruction. Specifically, podocytes were seeded on XFp microplates 24h before the experiment. On the day of the assay cells were rinsed and XF assay buffer was added for further equilibration. Afterwards the plate was incubated for 1 h at 37°C in a non-CO2 incubator. All medium and solutions of mitochondrial complex inhibitors were adjusted to pH 7.4 prior each assay. Following four baseline measurements of OCR and ECAR, for Mito

Stress Test inhibitors of the respiratory chain were sequentially injected into each well. Three OCR and ECAR readings were taken after addition of each inhibitor and before automated injection of the subsequent inhibitor. Mitochondrial complex inhibitors, in order of injection, included oligomycin (1.5 μM) to inhibit complex V, FCCP (1.0 μM) to uncouple the proton gradient, antimycin A (1.0 μM, inhibitor of complex III), and rotenone (1.0 μM, complex I inhibitor). OCR and ECAR were automatically calculated by Seahorse XFp software version 2.2.0 (Seahorse Bioscience, Billerica, MA, USA). After each experiment podocytes were fixed with paraformaldehyde and nuclei were stained with DAPI. Olympus ScanR Screening Station for high-throughput microscopy detection (Olympus, Tokyo, Japan) was used for assessment of cell number to normalize XFp analysis data.

*SNCA knockdown and over-expression in immortalized human podocytes*

Generation of siRNA mediated knockdown of SNCA in immortalized human podocytes was performed using the following sequences: T1-1: 5′-CAUAGUCAUUUCUAAAAGUUU-3′ T2-1: 5′- GGAUUUAUGUGGAUACAAAUU -3′. These sequences have been previously tested and published (20). Transfections were performed using Amaxa nucleofector technology (Lonza, Basel, Switzerland) according to manufacturer´s instructions. Cells were let 10 days to differentiate prior transfection and seeded back for 48 or 72 hours prior analysis. The knockdown efficiency was compared to scramble transfection using western blot 48 and 72 hours post transfection.

*Western blot and immunofluorescence*

The buffers, the systems and the protocols that were applied in western blotting can be found in our previous publication (21). The following anti-bodies were used in WB experiments: α-Galactosidase (OriGene Technologies), α-tubulin (Sigma-Aldrich), Actinin 4 (Abcam), p62 (Cell Signaling Technology), LC3 (Invitrogen), SNCA (Santa Cruz Biotechnology), SCARB2 (Lifespan Biosciences), TTYH3 (Origene), MFEG8 (Sigma-Aldrich), CD63 (Proteintech), DDP (Novus Biologicals), PLSCR3, GBA and ITM2B (Abcam). TOM20 (Santa Cruz Biotechnology) and LAMP-I (Abcam) were used for immunofluorescence staining. Immunofluorescence staining was performed on differentiated cells seeded in collagen IV 8-well chamber slides (Ibidi, Germany). Cells were fixed in 4% paraformaldehyde in PBS for 10 minutes and permeabilized using 0.1% Triton X-100 in PBS. Permeabilized cells were washed in PBS and blocked with 5% BSA in PBS for 1h at room temperature. Primary and secondary antibodies were diluted in blocking solution and incubated for 120 minutes and 45 minutes respectively.

*Immunostaining of SNCA*

Kidney biopsies before the onset of ERT and after five years of treatment were taken from a selection of a larger patient series already described elsewhere (22). Ethical permission was granted from the ethics committee of the Western Regional Health Authority in Norway (REK Vest no. 2010/2483). Informed consent was signed by the patient and/or their designees in all cases. Kidney biopsies from the Norwegian Kidney Biopsy Registry with normal light- or electron-microscopical appearance served as controls (REK Vest no. 2013/553). Paraffin sections of 3um thickness were cut and incubated at 42°C overnight. Sections were then deparaffinized and rehydrated (20 min xylene, 10 min 100% ethanol, and 5 min in 95%, 85%, 75% and 50% ethanol). Slides were washed for 5min in (PBS1X-tween0, 1%). Antigen Retrieval Solution (PH6) was used in steam cooker for 30 min and the slides were let to cool for 20 min on ice. Slides were washed 3 times with PBS and blocked with 5% BSA in PBS for 45 min. Endogenous peroxidase was inactivated with DAKO peroxidase inhibitor for 15 min. Primary antibodies were diluted in blocking solution (1:200) and the samples were incubated for 2 hours at room temperature. Goat anti-Rabbit Biotinylated - DAKO (1:500) was added for 30 min and Streptavidin\HRP- DAKO (1:500) for 30 min followed by ACE substrate\ DAKO for 10min. Slides were washed after every step with PBS1X. Finally the Slides were mounted using permanent mounting medium. Stained slides were scanned with the Aperio ScanScope XT system (Leica Biosystems Imaging, Wetzlar, Germany) at ×40 objective magnification and viewed in eSlide Manager (Leica Biosystems Imaging, Wetzlar, Germany). Staining intensities of SNCA were analyzed with the color deconvolution method(23). Percent total positive pixel count was acquired with the Aperio Color Deconvolution algorithm v9 (Leica Biosystems Imaging, Wetzlar, Germany) from annotated glomerular regions.

*In-cell western for SNCA*

Primary urinary cells were grown in a 0,1% Gelatine (Millipore Sigma) coated 96well plate until confluent. After treatment with Vehicle / lyso-Gb3, cells were washed in PBS, fixed in 4% Paraformaledhyde, washed again in PBS and blocked/permeabilized in 5% BSA, 0,1% Triton X in PBS for 1h at roomtemperature. After 3 washed in PBS, cells were stained with primary antibody (Sigma-Aldrich - HPA005459) 1:200 over night. The next day, after 3 additional washing steps in PBS, cells were incubated in secondary antibody (LI-COR IRDye 800cw) 1:200 and Draq5 nuclear stain (Abcam – 108410) 1:1000 or 1h at room temperature. Cells were washed again 3 times and imaged at the LI-COR Odyssey imager at 3µm focus, 89µm resolution.

*Statistics and reproducibility*

Data are expressed as mean ± SEM. Paired Student´s t-test or ONE-WAY Anova (multiple comparison test - Tukey) were used based on data distribution. Statistical significance was defined as *P<0.05, **P<0.01, ***P<0.001 and ****P<0.0001, n.s. - not statistically significant. Number of independent experiments and total amount of analysed cells are specified in each figure.

### Funding

FB was supported by the Else Kröner Fresenius Stiftung (2021_EKMS.26). WR acknowledges support by a starting grant from the European Research Council (Programme "Ideas," ERC-2011-StG 282105), and by the Deutsche Forschungsgemeinschaft (DFG, German Research Foundation) under Germany's Excellence Strategy (EXC-294 and EXC-2189). WR and CS were supported by the Freiburg Institute for Advanced Studies (FRIAS). VGP was supported by the DFG (CRC1192), by the BMBF (Fibromap), and the German Society of Nephrology DGFN. VGP was supported by the DFG (CRC1192), by the BMBF (Fibromap), and the German Society of Nephrology DGFN. CS was supported by the Excellence Initiative of the German Federal and State Governments, the German Society of Nephrology DGFN, the Else Kröner Fresenius Stiftung, NAKSYS and by the Berta-Ottenstein Programme, Faculty of Medicine, University of Freiburg. TBH was supported by the DFG (CRC1192, HU 1016/8-2, HU 1016/11-1, HU 1016/ 12-1), by the BMBF (STOP-FSGS- 01GM1901C, NephrESA-031L0191E and DEFEAT PANDEMIcs), by the Else-Kröner Fresenius Foundation (Else Kröner-Promotionskolleg –iPRIME), by the European Research Council-ERC (grant

616891) and by the H2020-IMI2 consortium BEAt-DKD (115974); this joint undertaking receives support from the European Union's Horizon 2020 research and innovation program and EFPIA and JDRF.

### Supplemental Figures

**Figure S1:**
(A) Two gRNAs were used to target the first exon within the *GLA* locus inducing premature stop codons. (B) Thin layer chromatography reveals a substantial accumulation of GB3 in the KO clones A and B. (C, D and E respectively) Mass spectrometry analysis of GB3 isoforms shows a significant accumulation of GB3-C24-1, GB3-C16-0 and GB3-C18-0 in GLA KO compared to WT cells. The expression level of these isoforms is nearly normalized upon 96hours of enzyme therapy.

**Figure S2:** (A) Seahorse XFp experiments confirms a normal mitochondrial function in KO cells (N=8). (B) Mitochondrial import receptor subunit (TOM20) staining in WT and KO cells is equally abundant and normally distributed. Scalebars indicating 10µm. (C) TEM images confirm a normal mitochondrial ultrastructure in KO cells. Scalebar indicating 500nm.

**Figure S3:**
(A) Western blot shows a comparable LC3-II accumulation and P62 level in GLA KO and WT cells after 2 hours of (100µM) chloroquine treatment. (B) Bands Quantification confirms a significant increase in LC3 in both clones compared to untreated cells but no difference between the genotypes has been noticed. (C) p62 in KO cells is less abundant in KO clones compared to WT cells at basal condition but not different upon chloroquine treatment (N=4).

**Figure S4:**
(A) RNA seq of WT and GLA-KO cells where GLA gene was the top downregulated gene (arrow). (B) Determining the module size of upregulated seed proteins. The z-score of the resulting module is plotted against the module size, which is determined by the known protein interactions of the DIAMOnD proteins and upregulated seed proteins with GLA. The red dashed line indicates the module size (number of genes n=64) where all upregulated seed proteins are integrated into the module. The resulting module has a significantly larger size than expected by random (z-score = 28). (C) Determining the module size of downregulated seed proteins. The z-score of the resulting module size is plotted against the module size, which is determined by the known protein interactions of the DIAMOnD proteins and downregulated seed proteins. The red dashed line indicates the module size (number of genes n=1185) where all downregulated seed proteins are integrated into the module. The resulting module has a significantly larger size than expected by random (z-score = 7).

**Figure S5:**
Overrepresented terms for the upregulated seed protein network (red) and downregulated seed protein network (blue) through (A) GO: cellular compartment, (B) Reactome pathway and (C) GO: molecular function annotation. FDR values were retrieved using the panther webtool (http://www.pantherdb.org, as of May 2019).

**Figure S6:**
Western blots confirming the over-expression of the top ten up regulated proteins (other than SNCA) which were reversible upon 96hours of aGAL therapy.

**Figure S7:**
(A & B) The quantification of in-cell western blot of SNCA vs Draq5 showing no significant differences between groups or upon Gb3 treatment (n=3 controls and 4 Fabry).

**Figure S8:**
(A) Western blots of SNCA and ACTN4 in WT vehicle-treated cells, KO vehicle-treated cells, KO 10µMClenbuterol-treated cells and KO cells treated with 20µMClenbuterol (B) The quantification of western blot bands of SNCA vs ACTNn4 showing a Clenbuterol dose dependent inhibition of SNCA (10 and 20 µM for 96 hours, N=6 per group).

**Figure S9**
(A) LAMP-1 staining in WT (D1), KO-A, KO-A treated with 20µM Clenbuterol and KO-A treated with 10 µM Orciprenaline shows a partial restoration of subcellular lysosomal distribution in beta 2 agonist treated KO-A cells compared to untreated ones. Scalebars indicating 10µm. (B) The quantitative ratiometric LysoSensor yellow/blue DND-160 assay demonstrates the beneficial significant effect of β2 adrenergic receptor agonists on Lysosomal pH (N=8). (C) Representative TEM images showing the ultrastructural change in the different treatment groups. Wild type cells treated with PBS. GLA knock out cells treated with PBS demonstrating drastic accumulation of multilaminar vacuoles. aGLA-KO cells treated with aGAL showing a restoration of the cellular ultrastructure. KO cells treated with 20µM Clenbuterol present a moderate decrease in the multilaminar vacuoles KO cells treated with recombinant aGAL and Clenbuterol showing a full restoration of the cellular ultrastructure. Cells in all groups were differentiated over 6 days and treated afterwards over 96 hours. Scalebars indicating 3µm and 1µm in detail.

### Supplemental Tables

**Table S1:** Patient characteristics
**Table S2:** Connectivity mapping analysis

### References:


1. Najafian B et al. One Year of Enzyme Replacement Therapy Reduces Globotriaosylceramide Inclusions in Podocytes in Male Adult Patients with Fabry Disease.. *PloS one* 2016;11(4):e0152812.

2. Najafian B et al. Accumulation of Globotriaosylceramide in Podocytes in Fabry Nephropathy Is Associated with Progressive Podocyte Loss.. *J Am Soc Nephrol Jasn* 2020;ASN.2019050497.

3. Najafian B et al. Progressive podocyte injury and globotriaosylceramide (GL-3) accumulation in young patients with Fabry disease. *Kidney international* 2010;79(6):663–670.

4. Toyoda M, Najafian B, Kim Y, Caramori ML, Mauer M. Podocyte Detachment and Reduced Glomerular Capillary Endothelial Fenestration in Human Type 1 Diabetic Nephropathy. *Diabetes* 2007;56(8):2155–2160.

5. Zhou T et al. Generation of human induced pluripotent stem cells from urine samples.. *Nature protocols* 2012;7(12):2080–2089.

6. Slaats GG et al. Urine-derived cells: a promising diagnostic tool in Fabry disease patients. *Sci Rep-uk* 2018;8(1):11042.

7. Yasuda-Yamahara M et al. FERMT2 links cortical actin structures, plasma membrane tension and focal adhesion function to stabilize podocyte morphology.. *Matrix biology : journal of the International Society for Matrix Biology* 2018;68–69:263–279.

8. Grahammer F et al. A flexible, multilayered protein scaffold maintains the slit in between glomerular podocytes. *Jci Insight* 2016;1(9):e86177.

9. Subramanian A et al. A Next Generation Connectivity Map: L1000 Platform and the First 1,000,000 Profiles. *Cell* 2017;171(6):1437-1452.e17.

10. Cheng F et al. Network-based approach to prediction and population-based validation of in silico drug repurposing. *Nat Commun* 2018;9(1):2691.

11. Ghiassian SD, Menche J, Barabási A-L. A DIseAse MOdule Detection (DIAMOnD) algorithm derived from a systematic analysis of connectivity patterns of disease proteins in the human interactome.. *Plos Comput Biol* 2015;11(4):e1004120.

12. Halu A et al. Exploring the cross-phenotype network region of disease modules reveals concordant and discordant pathways between chronic obstructive pulmonary disease and idiopathic pulmonary fibrosis.. *Hum Mol Genet* 2019;28(14):2352–2364.

13. Beutler E, Kuhl W. Purification and properties of human alpha-galactosidases.. *J Biological Chem* 1972;247(22):7195–200.

14. Kint JA. Fabry's Disease: Alpha-Galactosidase Deficiency. *Science* 1970;167(3922):1268–1269.

15. Mayes JS, Scheerer JB, Sifers RN, Donaldson ML. Differential assay for lysosomal alpha-galactosidases in human tissues and its application to Fabry's disease. *Clin Chim Acta* 1981;112(2):247–251.

16. Shen J-S et al. Mannose receptor-mediated delivery of moss-made α-galactosidase A efficiently corrects enzyme deficiency in Fabry mice.. *J Inherit Metab Dis* 2015;39(2):293–303.

17. Luciani A et al. Impaired Lysosomal Function Underlies Monoclonal Light Chain-Associated Renal Fanconi Syndrome.. *J Am Soc Nephrol Jasn* 2015;27(7):2049–61.

18. Kubota C et al. Constitutive reactive oxygen species generation from autophagosome/lysosome in neuronal oxidative toxicity.. *J Biological Chem* 2009;285(1):667–74.

19. Bligh EG, Dyer WJ. A RAPID METHOD OF TOTAL LIPID EXTRACTION AND PURIFICATION. *Can J Biochem Phys* 1959;37(8):911–917.

20. Takahashi M et al. Normalization of Overexpressed α-Synuclein Causing Parkinson's Disease By a Moderate Gene Silencing With RNA Interference.. *Mol Ther Nucleic Acids* 2015;4(Lancet 373 2009):e241.

21. Rogg M et al. The WD40-domain containing protein CORO2B is specifically enriched in glomerular podocytes and regulates the ventral actin cytoskeleton.. *Scientific Reports* 2017;7(1):15910.

22. Skrunes R et al. Long-Term Dose-Dependent Agalsidase Effects on Kidney Histology in Fabry Disease.. *Clinical journal of the American Society of Nephrology : CJASN* 2017;12(9):1470–1479.

23. Ruifrok AC, Johnston DA. Quantification of histochemical staining by color deconvolution.. *Anal Quantitative Cytol Histology Int Acad Cytol Am Soc Cytol* 2001;23(4):291–9.
