## Supplementary figures and images for "Synuclein alpha accumulation mediates podocyte injury in Fabry nephropathy"

### Supplementary Figures

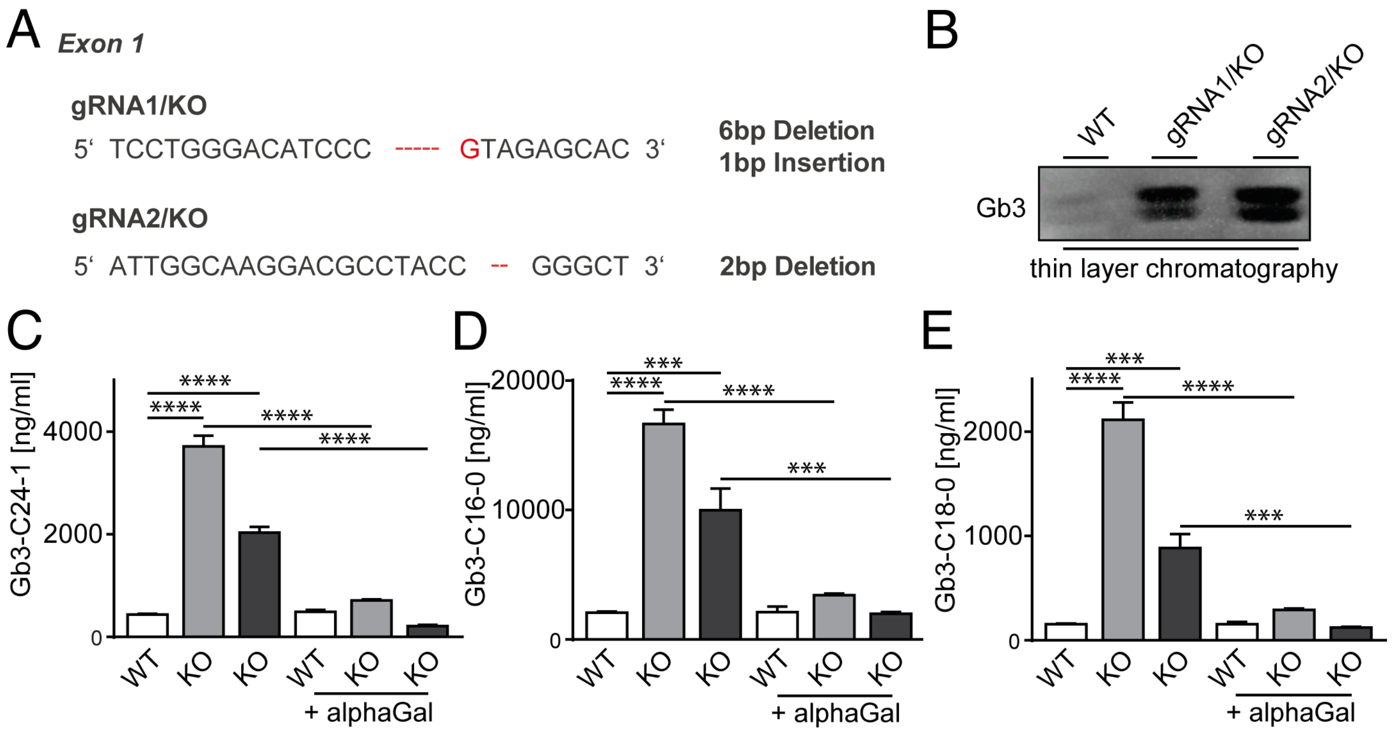

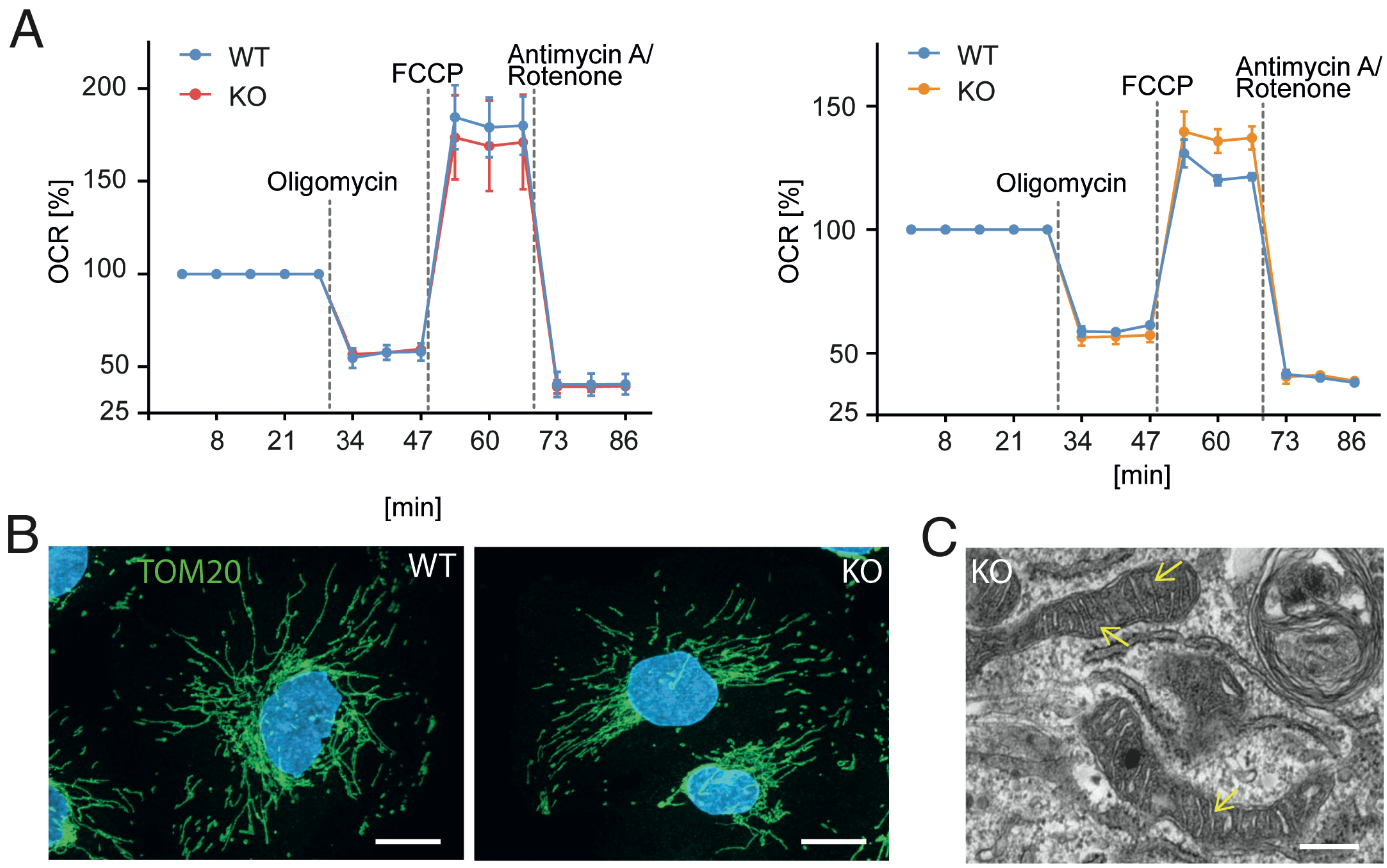

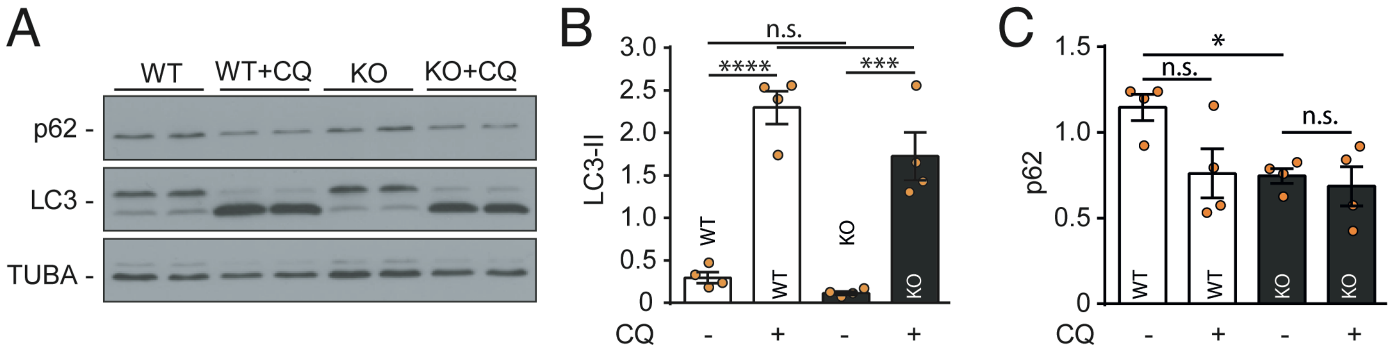

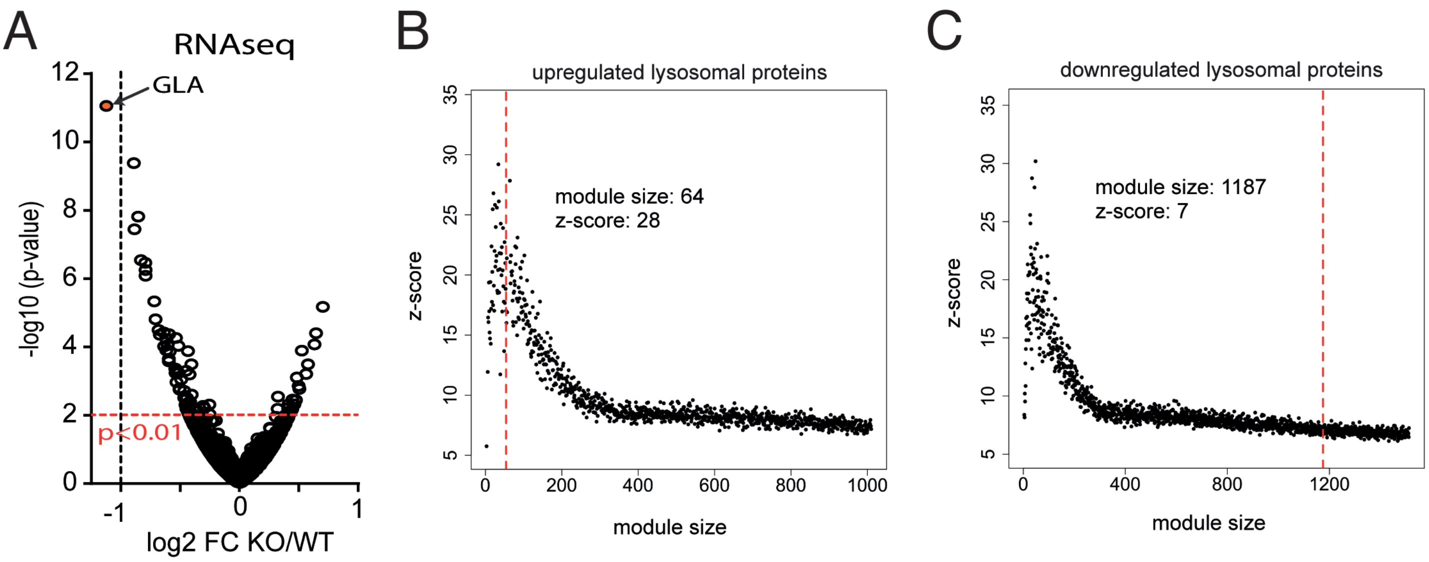

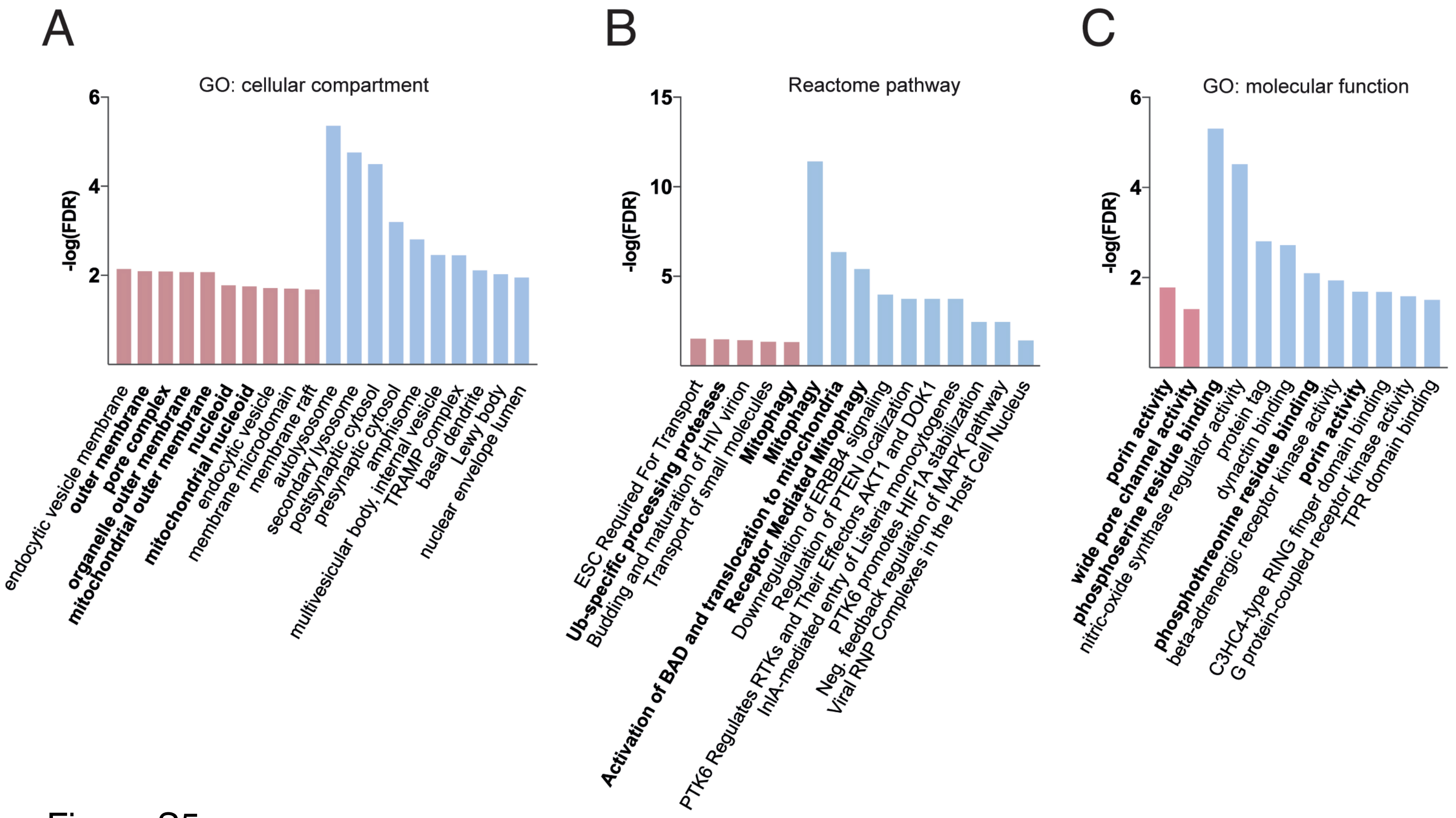

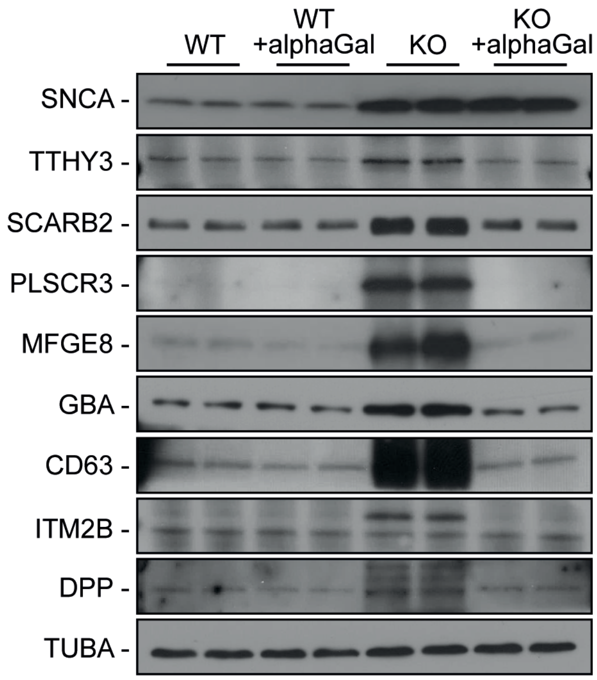

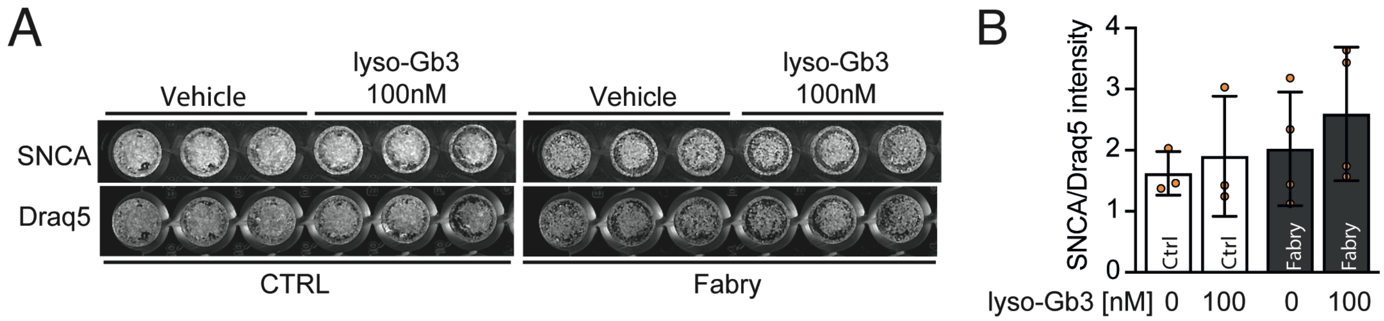

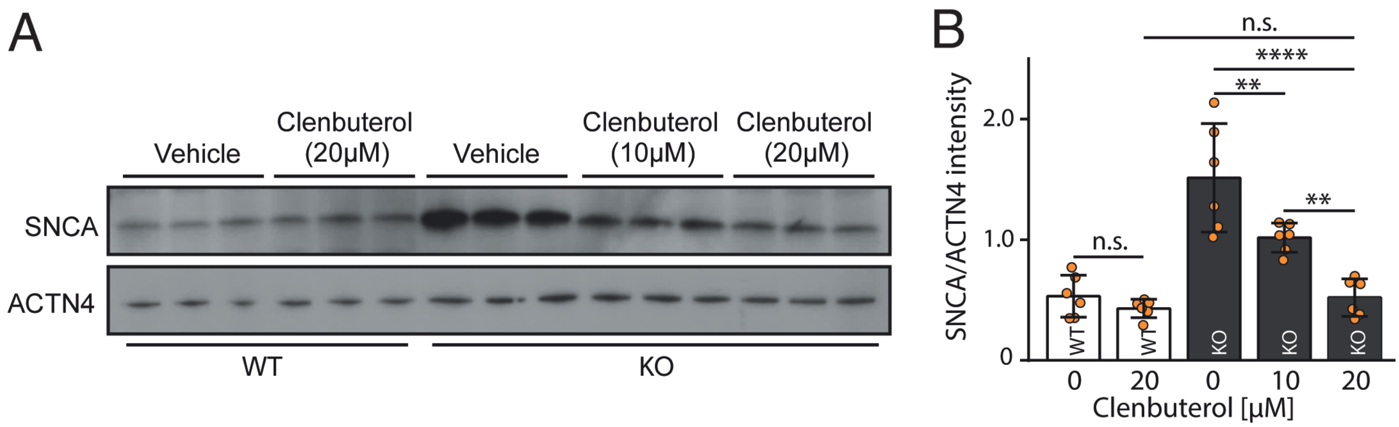

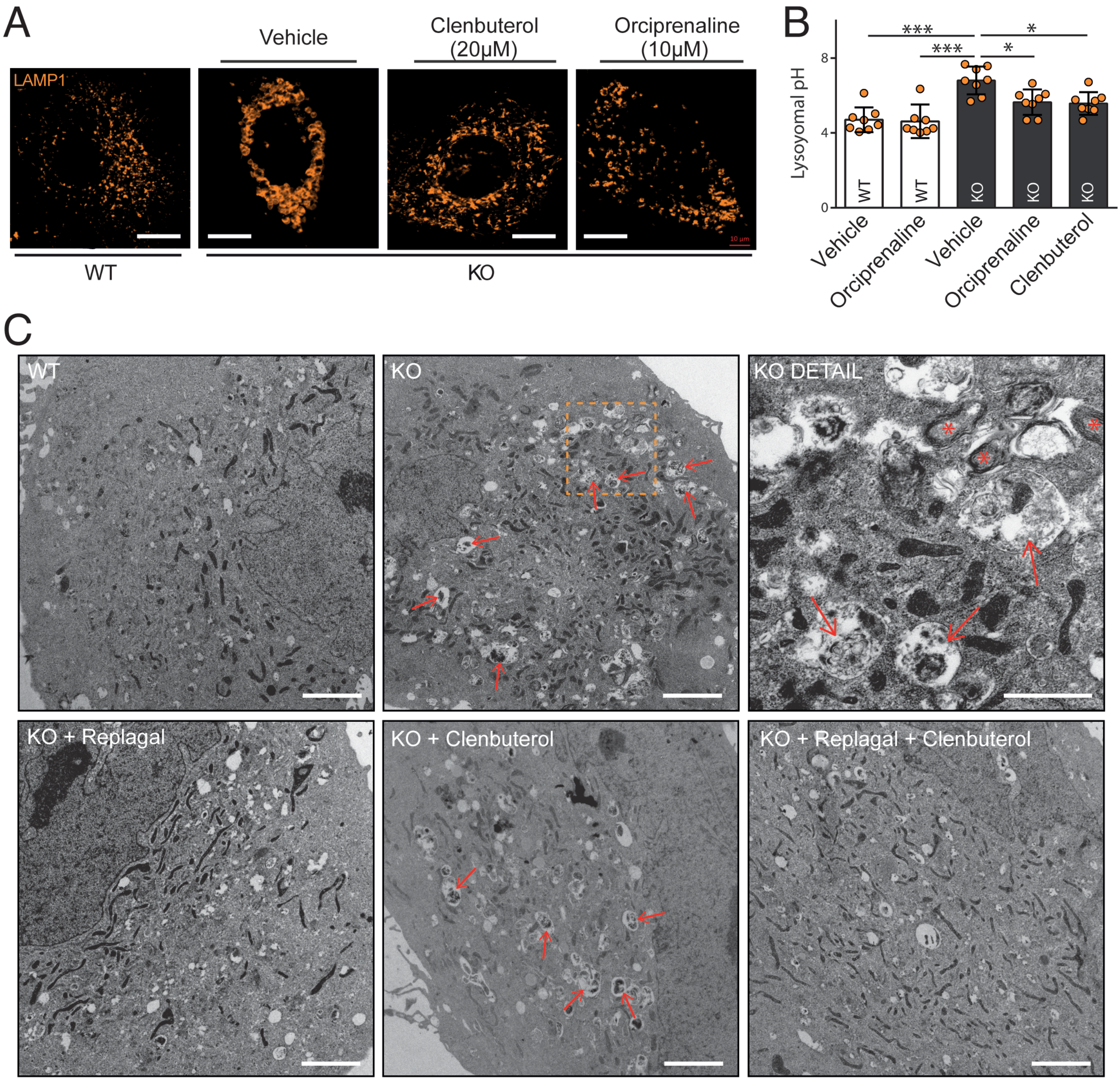
